# Unexpectedly low recombination rates and presence of hotspots in termite genomes

**DOI:** 10.1101/2024.03.22.586269

**Authors:** Turid Everitt, Tilman Rönneburg, Daniel Elsner, Anna Olsson, Yuanzhen Liu, Tuuli Larva, Judith Korb, Matthew T Webster

## Abstract

Meiotic recombination is a fundamental evolutionary process that facilitates adaptation and the removal of deleterious genetic variation. Social Hymenoptera exhibit some of the highest recombination rates among metazoans, whereas high recombination rates have not been found among non-social species from this insect order. It is unknown whether elevated recombination rates are a ubiquitous feature of all social insects. In many metazoan taxa, recombination is mainly restricted to hotspots a few kilobases in length. However, little is known about the prevalence of recombination hotspots in insect genomes. Here we infer recombination rate and its fine-scale variation across the genomes of two social species from the insect order Blattodea: the termites *Macrotermes bellicosus* and *Cryptotermes secundus*. We used linkage-disequilibrium-based methods to infer recombination rate. We infer that recombination rates are close to 1 cM/Mb in both species, similar to the average metazoan rate. We also observed a highly punctate distribution of recombination in both termite genomes, indicative of the presence of recombination hotspots. We infer the presence of full-length *PRDM9* genes in the genomes of both species, which suggests recombination hotspots in termites might be determined by *PRDM9*, as they are in mammals. We also find that recombination rates in genes are correlated with inferred levels of germline DNA methylation. The finding of low recombination rates in termites indicates that eusociality is not universally connected to elevated recombination rate. We speculate that the elevated recombination rates in social Hymenoptera are instead promoted by intense selection among haploid males.

## Introduction

Meiotic recombination has two fundamental roles: It is essential for correct disjunction of chromosomes during cell division and it generates new combinations of genetic variants that form the raw material of evolution (Hartfield and Keightley 2012). However, recombination can also be deleterious, as it can generate structural mutations and break up favourable combinations of alleles. Recombination varies among species, with optimal recombination rate in a genome likely determined by the optimal trade-off of its positive and negative evolutionary and mechanistic effects (Stapley et al. 2017).

The highest recombination rates in metazoans observed so far are found in social insects (Wilfert et al. 2007; Stapley et al. 2017). However, until now recombination rate has only been studied in social insects from the order Hymenoptera, to which the majority of social insects belong. Among Hymenoptera, the highest rates are found in the genus *Apis*: the Western honey bee *Apis mellifera* (20.8 cM/Mb) and its relatives *Apis cerana* (17.4 cM/Mb), *Apis florea* (20.8 cM/Mb) and *Apis dorsata* (25.1 cM/Mb) (Beye et al. 2006; Shi et al. 2013; Liu et al. 2015; Wallberg et al. 2015; Rueppell et al. 2016; Kawakami et al. 2019). Other social Hymenoptera also have high rates including the bumblebee *Bombus terrestris* (8.9 cM/Mb) (Liu et al. 2017; Kawakami et al. 2019), the stingless bee *Frieseomelitta varia* (9.3 – 12.5 cM/Mb) (Waiker et al. 2021), the wasp *Vespula vulgaris* (9.7 cM/Mb) (Sirviö et al. 2011a) and the ants *Pogonomyrmex rugosus* (11.1 cM/Mb) (Sirviö et al. 2011b) and *Acromymex echinatior* (6.1 cM/Mb) (Sirviö et al. 2006). In contrast, the solitary bee *Megachile rotundata* and the solitary wasp *Nasonia* have relatively low rates (1.0 and 1.5 cM/Mb, respectively) (Niehuis et al. 2010; Jones et al. 2019). For comparison, the average recombination rate of 15 insect species outside of Hymenoptera is 2.2 cM/Mb (Wilfert et al. 2007).

We can learn about the forces shaping the evolution of recombination rates in general by understanding why high recombination rates have evolved in social Hymenoptera. Several hypotheses have been proposed to explain this observation. One set of explanations focusses on the effects of recombination on intra-colony genetic diversity. For example, increasing the genetic diversity of a colony could promote task specialization among workers (Kent et al. 2012; Kent and Zayed 2013). Similarly, elevated genetic variation in a colony could prevent invasion by pathogens or parasites due to diversification of immune genes (Fischer and Schmid-Hempel 2005). Recombination could also reduce variance in relatedness between nestmates, thereby reducing potential kin conflict within colonies (Sherman 1979; Templeton 1979; Wilfert et al. 2007). Additionally, it has been proposed that high recombination rates could have facilitated the evolution of eusociality over a longer evolutionary timescale, because recombination between genes that take on functions in different castes permits them to evolve more independently (Kent and Zayed 2013). Several studies have addressed these hypotheses (Kent et al. 2012; Liu et al. 2015; Wallberg et al. 2015; Rueppell et al. 2016; Liu et al. 2017; Jones et al. 2019; Waiker et al. 2021; Kawakami et al. 2019) but so far none is strongly supported. It is also not clear whether eusociality *per se* selects for higher recombination rates, or whether high rates are a specific feature of eusocial Hymenoptera.

The main characteristics of eusocial insect species are the presence of castes that forgo reproduction in order to care for brood or defend other colony members (workers and soldiers). Eusocial insects are found mainly in Hymenoptera, among bees, wasps and ants, and also Blattodea, in which all termites (Infraorder: Isoptera) are eusocial but exhibit varying levels of social complexity. These insect orders likely diverged ∼400 million years ago (Misof et al. 2014) and there are substantial differences between social insects from the two orders (Korb 2008; Korb and Thorne 2017). Hymenoptera belong to the superorder Holometabola, which is the most diverse insect superorder and contains 11 orders including Lepidoptera (butterflies, moths), Coleoptera (beetles), and Diptera (true flies), which all undergo complete metamorphosis. Blattodea is a hemimetabolous order, which undergoes partial metamorphosis. Hymenoptera have haplodiploid sex determination in which males develop from haploid, unfertilized eggs and females derive from diploid, fertilized eggs. In Blattodea, both males and females are diploid. Members of the worker caste in eusocial Hymenoptera are all female, whereas eusocial Blattodea workers are of both sexes. Hymenoptera colonies are headed by one or a small number of queens, whereas Blattodea colonies contain both a king and a queen.

The hypotheses proposed to explain the high recombination rates observed in eusocial Hymenoptera predict that recombination should be elevated in all social insects. However, so far it is unknown whether social insects from other orders also have high rates. As described above, there are fundamental differences in the sex determination system and colony compositions of social insects from Hymenoptera and Blattodea. The robustness of the association between eusociality and high recombination rates can therefore be addressed by assessing mean genomic recombination rates in social Blattodea (termites).

In addition to variation between species, the rate of recombination also varies along chromosomes. In mammals, crossover events are mainly restricted to regions that contain specific motifs that act as binding sites for the PRDM9 (PR/SET domain 9) protein that marks regions for double-stranded breaks, which are repaired by recombination during meiosis (Baudat et al. 2010; Myers et al. 2010; Parvanov et al. 2010). Species with an active PRDM9 protein include nearly all mammals and many vertebrate taxa (Baker et al. 2017). In species without PRDM9, recombination can be directed to hotspots defined by other features, such as CpG islands and promoters that have open chromatin. Such hotspots have been characterised in the genomes of diverse taxa including yeast, birds, and dogs (Axelsson et al. 2012; Lam and Keeney 2015; Singhal et al. 2015; Berglund et al. 2014). In insects, the prevalence of recombination hotspots is not well studied. Fine-scale recombination maps in the fruit fly *Drosophila melanogaster* and honey bee *A. mellifera* indicate a paucity of recombination hotspots (Chan et al. 2012; Smukowski Heil et al. 2015; Wallberg et al. 2015), whereas hotspots appear to be present in the genome of the butterfly *Leptidea sinapis* (Torres et al. 2023).

Several other factors may also modulate recombination rate along chromosomes, such as germline DNA methylation. Many vertebrate species have elevated recombination in CpG islands, which are usually unmethylated in the germline (Axelsson et al. 2012; Singhal et al. 2015; Berglund et al. 2014). In insect genomes, methylation is not always present, but commonly restricted to gene bodies (Arsala et al. 2022; Wang et al. 2013; Lyko et al. 2010; Ventós-Alfonso et al. 2020; Bewick et al. 2019; Harrison et al. 2018). In the honey bee *A. mellifera* and solitary bee *M. rotundata,* recombination is reduced in genes, particularly those inferred to have the highest levels of germline methylation (Wallberg et al. 2015; Jones et al. 2019). In addition, some studies have identified correlations between caste-biases in gene expression and recombination. In honey bees, genes with worker-biased gene expression have been found to have elevated recombination rates (Kent et al. 2012) but this is likely an indirect result of differences in germline methylation between sets of genes (Wallberg et al. 2015).

Here we estimate recombination rate and its fine-scale variation across the genomes of two distantly related termite species with contrasting social complexities (Korb and Hartfelder 2008; Korb and Thorne 2017). *Macrotermes bellicosus* (Termitidae: Macrotermitinae) is a fungus-growing termite with a high level of social complexity, with colonies of several millions of individuals and sterile workers. It belongs to the foraging (multiple-pieces nesting) termites, where workers leave the nest to forage for food. The drywood termite *Cryptotermes secundus* (Kalotermitidae) has a low level of social complexity. Its colonies consist of a few hundred individuals and totipotent workers from which queens and kings develop (Hoffmann et al. 2012). It belongs to the wood-dwelling (one-piece nesting) termites that nest in a single piece of wood that also serves as their food source.

The study has three main aims. Firstly, we aim to test the robustness of the association between sociality and high recombination rate. If sociality is a universal driver of high recombination rates, we would expect both termite species to have elevated recombination rates, and that it is particularly elevated in *M. bellicosus*, which has a higher level of social complexity. Secondly, we aim to determine whether recombination hotspots exist in the two termite genomes, and infer whether active full-length *PRDM9* genes are present in their genomes. Thirdly, we aim to analyse the factors that govern recombination rate variation across the genome, in particular whether it is associated with patterns of methylation and gene expression as has been reported in other studies.

## Results

### Genetic variation in two termite species is typical for social insects

We utilized genome assemblies of two termite species: *M. bellicosus* and *C. secundus*. The genome assembly of *M. bellicosus* (Qiu et al. 2023) is 1.3 Gbp in total length across 275 scaffolds with scaffold N50 of 22.4 Mbp. The genome assembly of *C. secundus* (Csec_1.0) (Harrison et al. 2018) is divided into 55,483 scaffolds with a scaffold N50 of 1.2 Mbp and a total length of 1.0 Gbp.

We sequenced 10 unrelated individuals each of *M. bellicosus* and *C. secundus* to a mean depth of 34x and 31x per sample, respectively (Supplemental Table S1). We mapped reads to the appropriate genome assemblies and called variants in each species. In *M. bellicosus* we called 5.6 million SNPs and in *C. secundus* 15.1 million SNPs (Table 1). One individual of *C. secundus* (CS_8) was excluded due to a low proportion of mapped reads, poor mapping quality and strand bias in mapping. Estimates of variation based on *θ*_*W*_ per bp (Watterson 1975) are 0.14% and 0.44% in *M. bellicosus* and *C. secundus* respectively, which are broadly comparable to levels of variation in the honey bee *A. mellifera* (0.3 - 0.8%) (Wallberg et al. 2014).

**Table 1.**
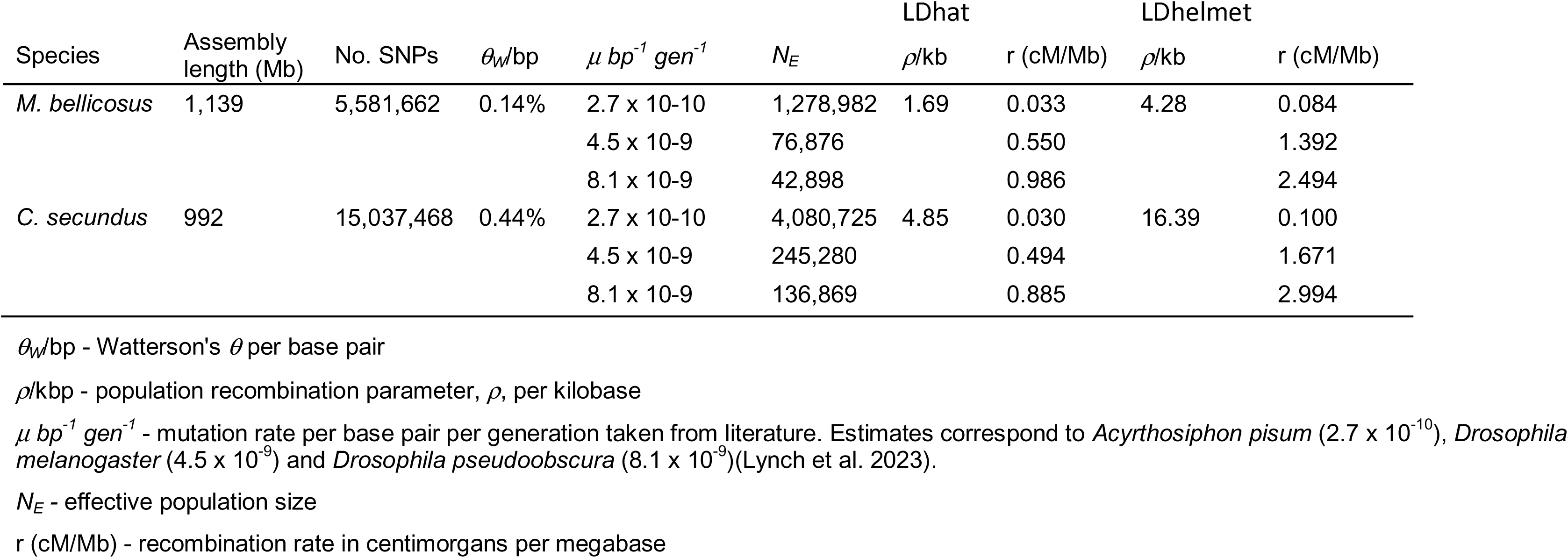
Estimates of levels genetic variation, effective population size and recombination rate in two termite species.

| Species | Assembly<br>length (Mb) | No. SNPs | $\theta_W/\text{bp}$ | $\mu \text{ bp}^{-1} \text{ gen}^{-1}$ | $N_E$ | LDhat | | LDhelmet | |
| --- | --- | --- | --- | --- | --- | --- | --- | --- | --- |
| | | | | | | $\rho/\text{kb}$ | $r \text{ (cM/Mb)}$ | $\rho/\text{kb}$ | $r \text{ (cM/Mb)}$ |
| *M. bellicosus* | 1,139 | 5,581,662 | 0.14% | 2.7 x 10-10 | 1,278,982 | 1.69 | 0.033 | 4.28 | 0.084 |
|  |  |  |  | 4.5 x 10-9 | 76,876 |  | 0.550 |  | 1.392 |
|  |  |  |  | 8.1 x 10-9 | 42,898 |  | 0.986 |  | 2.494 |
| *C. secundus* | 992 | 15,037,468 | 0.44% | 2.7 x 10-10 | 4,080,725 | 4.85 | 0.030 | 16.39 | 0.100 |
|  |  |  |  | 4.5 x 10-9 | 245,280 |  | 0.494 |  | 1.671 |
|  |  |  |  | 8.1 x 10-9 | 136,869 |  | 0.885 |  | 2.994 |

$\theta_W/\text{bp}$ - Watterson's $\theta$ per base pair $\rho/\text{kbp}$ - population recombination parameter, $\rho$ , per kilobase $\mu \text{ bp}^{-1} \text{ gen}^{-1}$ - mutation rate per base pair per generation taken from literature. Estimates correspond to *Acyrtosiphon pisum* ( $2.7 \times 10^{-10}$ ), *Drosophila* *melanogaster* ( $4.5 \times 10^{-9}$ ) and *Drosophila pseudoobscura* ( $8.1 \times 10^{-9}$ )(Lynch et al. 2023). $N_E$ - effective population size $r \text{ (cM/Mb)}$ - recombination rate in centimorgans per megabase

We used these estimates of genetic variation to estimate effective population size (*N*_*E*_) for each of the sampled populations (Table 1). These rely on an estimate of mutation rate, *μ*, in each species. As there is no estimate of *μ* for Blattodea, we considered a range of estimates from insects compiled by (Lynch et al. 2023) based on experimental assays that includes estimates from Diptera (*Drosophila*) (Keightley et al. 2009, 2014), Hymenoptera (Liu et al. 2017; Yang et al. 2015) and Lepidoptera (Keightley et al. 2015). These estimates range between a minimum of 2.7 x 10^-10^ bp^-1^gen^-1^ in *Acyrthosiphon pisum* (Fazalova and Nevado 2020) to a maximum of 8.1 x 10^-9^ bp^-1^gen^-1^ in *Drosophila pseudoobscura* (Krasovec 2021). The average of multiple estimates in *Drosophila melanogaster* is 4.5 x 10^-9^ bp^-1^gen^-1^, which is typical of insects (Lynch et al. 2023). A mutation rate of *μ* = 1.85 x 10^-9^ bp^-1^gen^-1^ has been estimated in the orchid mantis, *Hymenopus coronatus* (Huang et al. 2023) based on inference from levels of interspecific divergence. This species is a member of the order Mantodea, which contains the closest relatives of Blattodea.

Our estimates of *N*_*E*_ based on *θ*_*W*_ and the various mutation rates are ∼43,000 – 1,279,000 in *M. bellicosus*, which has high social complexity, and ∼137,000 – 4,081,000 in *C. secundus*, which has a lower level of social complexity (Table 1). Estimates of *N*_*E*_ assuming the typical mutation rate of 4.5 x 10^-9^ bp^-1^gen^-1^ are 77,000 and 245,000 respectively. Assuming the mutation rate of *H. coronatus* gives an *N*_*E*_ of 187,000 for *M. bellicosus* and 596,000 for *C. secundus*. These estimates are consistent with previous evidence that eusocial species with large colonies and high social complexity tend to have lower effective population sizes and the values are broadly comparable to levels of *N*_*E*_ found in social and solitary Hymenoptera (Leffler et al. 2012; Romiguier et al. 2014).

### Low average recombination rate in termite genomes

We next examined the decay of linkage disequilibrium (LD) based on the statistic *r*^2^. Both termite species show more extensive LD compared to *A. mellifera*, which has an extremely high recombination rate (Wallberg et al. 2015) (Figure 1). As *N*_*E*_ is broadly similar in all these species, these differences are expected to largely represent differences in recombination rates, and indicate that the two termite species do not exhibit the elevated genome average recombination rate that is observed in honey bees.

**Figure 1.**
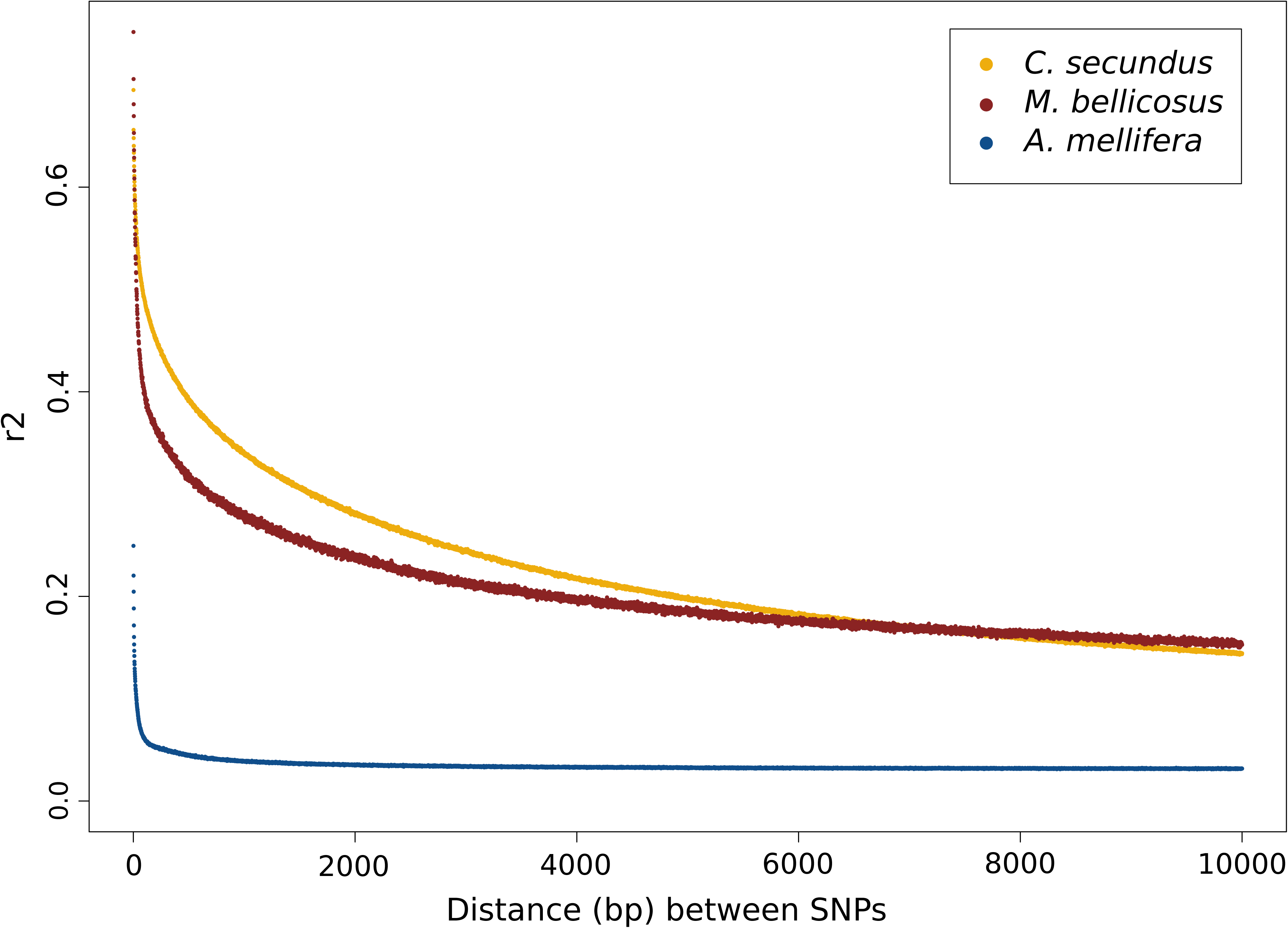
Genome-wide decay of LD measured as *r*^2^ versus distance (bp) between SNPs for the two termite species (*M. bellicosus*, dark red; *C. secundus*, yellow) and honey bees (*A. mellifera scutellata*, blue; data from Wallberg et al. 2017).

We estimated variation in the population recombination rate, *ρ*, in both datasets using LDhelmet (Chan et al. 2012) and LDhat (Auton and McVean 2007) (Table 1). Average rates of *ρ*/kbp in *M. bellicosus* and *C. secundus* are 1.7 and 4.8 respectively from LDhat and 4.28 and 16.39 respectively from LDhelmet. These estimates can be converted to cM/Mb using estimates of *N*_*E*_, which gives average recombination rates of 0.033 - 0.986 cM/Mb in *M. bellicosus* and 0.030 - 0.885 cM/Mb in *C. secundus* from LDhat and 0.084-2.49 cM/Mb for *M. bellicosus* and 0.1 - 2.99 cM/Mb for *C. secundus* from LDhelmet. Using a mutation rate close to the average for insects, the recombination rates in *M. bellicosus* and *C. secundus* are 0.550 and 0.494 cM/Mb respectively from LDhat and 1.39 and 1.67 cM/Mb respectively from LDhelmet. Using the mutation rate from *H. coronatus*, the recombination rates are 0.23 and 0.20 cM/Mb from LDhat and 0.57 and 0.69 cM/Mb from LDhelmet for *M. bellicosus* and *C. secundus*, respectively. Even though these estimates vary depending on the algorithm used for estimating *ρ* and the assumed mutation rate, they are all substantially lower than estimates from eusocial Hymenoptera (6-25 cM/Mb; (Sirviö et al. 2006) and more similar to rates found in other insects (Wilfert et al. 2007). This represents the first estimation of recombination rate in social insects outside of Hymenoptera and demonstrates that eusociality is not a universal driver of high recombination rates. Spearman’s correlation between the results from LDhat and LDhelmet, using 10 kbp windows, was *ρ* = 0.946 and *ρ* = 0.947 for *M. bellicosus* and *C. secundus*, respectively, with p < 2.2 x 10^-16^ for both species. For the remaining analyses, results from the software LDhelmet were used.

### Evidence for active recombination hotspots in termite genomes

Recombination rates are highly variable across the genomes of *M. bellicosus* (Figure 2A) and *C. secundus* (Figure 2B). We characterised the distribution of recombination events across the two termite genomes using a cumulative distribution plot (Figure 3). We found that 50% of the recombination occurs in 0.4% of the genome in *M. bellicosus* and 0.2% of the genome in *C. secundus*. This indicates that the majority of recombination events are restricted to a much smaller portion of the genome than observed in *A. mellifera*, where 50% of recombination events occur in 32% of the genome (Wallberg et al. 2015).

**Figure 2.**
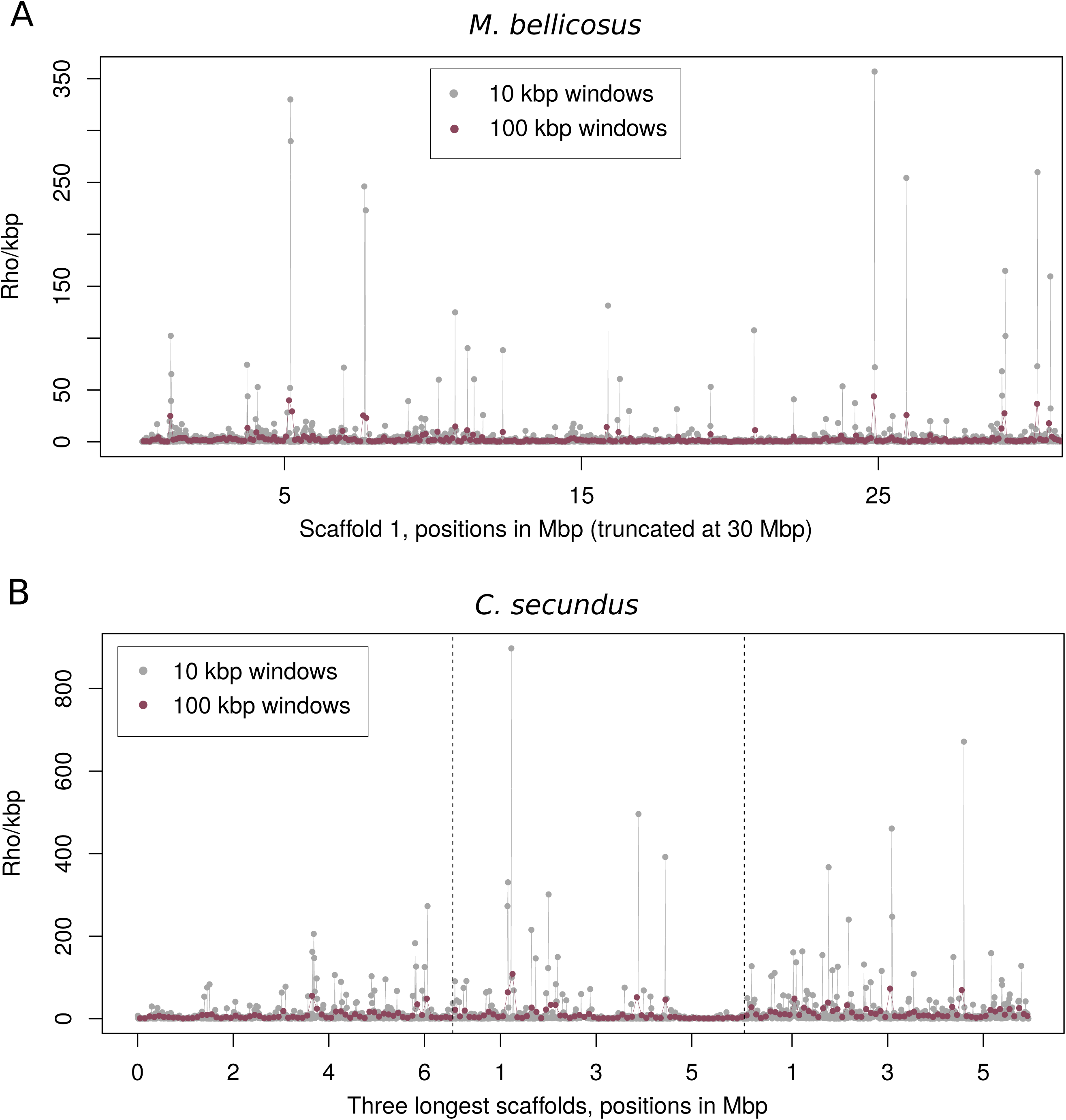
Variation in recombination rate estimated as *ρ*/kbp across representative genomic scaffolds in a) *M. bellicosus* and b) *C. secundus*. Estimates of *ρ*/kbp are calculated in 10 kbp and 100 kbp windows.

**Figure 3.**
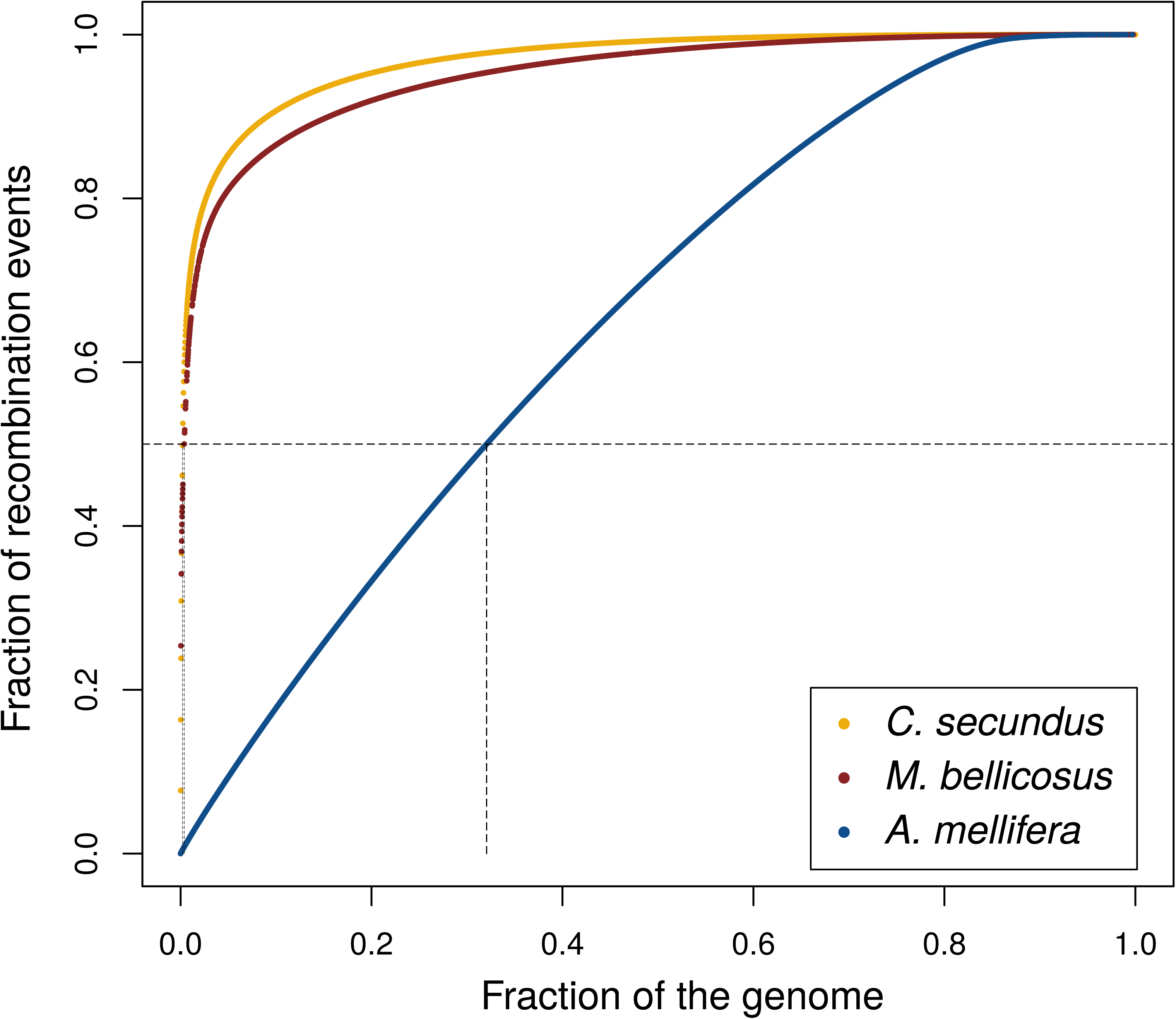
Cumulative plot of proportion of recombination events versus proportion of the genome for the two termite species *M. bellicosus* and *C. secundus* compared to the honey bee *A. mellifera* (Wallberg et al. 2015). Dashed lines show the proportion of the genome where 50 % of the recombination occurs, which for *M. bellicosus* is 0.4 %, *C. secundus* is 0.2 %, and *A. mellifera* is 32 %.

We further investigated whether recombination hotspots are present in the two termite genomes by looking at the distribution of *ρ* in windows of 1 kbp, 10 kbp and 100 kbp across the genome (Supplemental Figure S1). For all window sizes, a small subset of windows was observed with *ρ* greatly exceeding the mean in both termite species. For example, the percentage of 1 kbp windows with recombination rate at least four standard deviations above the genome wide mean value is 0.2% for *M. bellicosus* and 0.6% for *C. secundus*. By contrast, *A. mellifera* has no windows above this limit for any window size, consistent with a lack of recombination hotspots.

We next defined hotspots as 2 kbp regions with more than 5-fold higher recombination rate than the surrounding 100 kbp and used STREME (Bailey 2021) to identify 8 to 20-mers that were enriched in hotspots compared to the rest of the genome. This yielded significantly enriched motifs in both species (4 motifs for *M. bellicosus* and 1 motif for *C. secundus* after compensating for multiple hypothesis testing; Supplemental Tables S2 and S3). However, none of the motifs we identified are present in more than 5% of hotspots or exhibit more than 2-fold difference in frequency of occurrence relative to the background, suggesting they are not viable candidate motifs for promoting recombination in hotspots.

### Identification and characterization of the PRDM9 gene in termites

We searched the *M. bellicosus* and *C. secundus* genomes for evidence of complete *PRDM9* genes (Figure 4A), which could potentially govern the presence of recombination hotspots in these genomes, as it has been shown to do in mammalian genomes (Myers et al. 2010; Baudat et al. 2010; Parvanov et al. 2010). The *C. secundus* genome contains a *PRDM9* homolog (XP_023708049.2) with annotated KRAB, SSXRD, SET and zinc finger domains (Figure 4B). There is no complete homolog of *PRDM9* in the *M. bellicosus* annotation. We therefore used BLAST to search for sequences homologous to the conserved functional domains from *C. secundus*. We identified a putative *PRDM9* orthologue on Scaffold 42 of the *M. bellicosus* genome assembly, containing all domains in the correct order. We identified retroviral conserved domains in an intron 5’ of the zinc-finger domains in this gene, which are not predicted to alter the transcript. We confirmed the presence of a full-length transcript as well as presence and correct order of conserved domains by alignment of the *Zootermopsis nevadensis* PRDM9 protein sequence to the *M. bellicosus* genome using GeneWise (Madeira et al. 2024)(Figure 4C and Supplemental Table S4).

**Figure 4.**
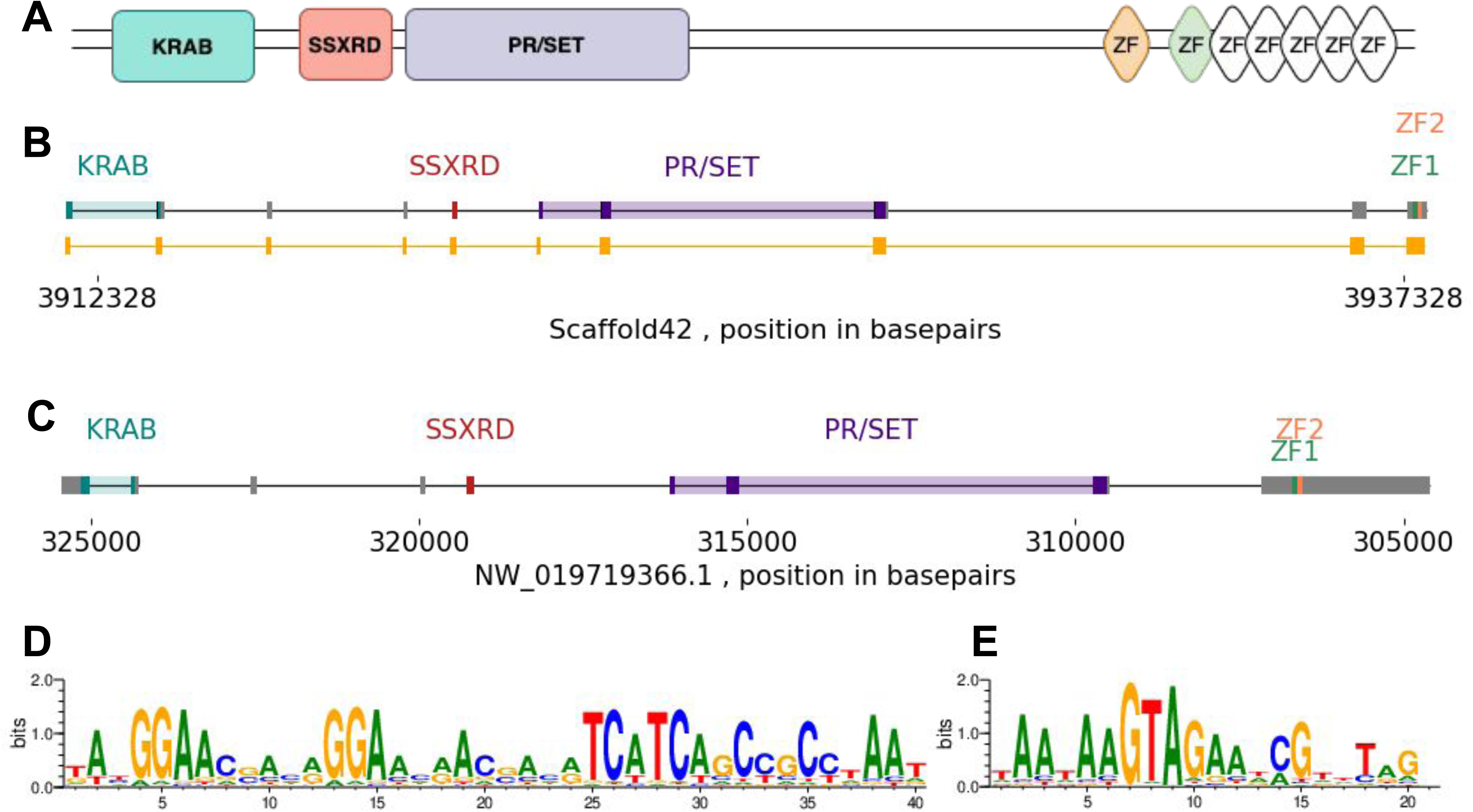
PRMD9 structure and predicted zinc-finger motifs in the *M. bellicosus* and *C. secundus* genomes. A) Predicted structure of the PRDM9 protein sequence in *C. secundus.* B) Predicted structure of the *PRDM9* gene in *M. bellicosus* identified by homology searches. Exons and conserved functional domains are marked. The mapped exons from *Zoothermopsis nevadensis* are also shown. C) Structure of *PRDM9* gene in *C. secundus*. Exons and conserved functional domains are marked. D) Predicted zinc finger binding motif in *M. bellicosus,* E) Predicted zinc finger binding motif in *C. secundus*.

We used a zinc-finger prediction algorithm (Persikov and Singh 2014) to predict DNA binding motifs from the zinc-finger domain present in PRDM9 in the *M. bellicosus*and *C. secundus* genome assemblies. The most probable consensus motifs were TATGGAACGACAGGAACAACGACATCATCAGCCGCCTAAT and TAATAAGTAGAATCGTTTAG for *M. bellicosus* (Figure 4D) and *C. secundus* (Figure 4E) respectively. Base probability matrices for these motifs are shown in Supplemental Table S5. We tested whether the presence of these motifs corresponded to elevated recombination considering 1 kbp around each motif. However, recombination rate was not significantly elevated around these predicted motifs for either species, either when searching based on the most likely motif, or by searching based on the probability matrices.

### Genomic correlates of recombination rate

We analysed genomic correlates of recombination rate in 10 kbp windows. There is a significant correlation between recombination and CpG_O/E_ in both species with Spearman’s *ρ* = 0.24, p < 1 x 10^-4^ for *M. bellicosus* and Spearman’s *ρ* = 0.10, p < 1 x 10^-4^ for *C. secundus* (Supplemental Figure S2). A reduction of recombination rates in regions of low CpG content is consistent with interference of germline methylation with recombination. However, as the correlation coefficients are low, the explanatory power is weak. Correlations of similar magnitudes were observed between recombination and GC content (Spearman’s *ρ* =0.23 for *M. bellicosus*, p < 1 x 10^-4^ and Spearman’s *ρ* =0.017 for *C. secundus*, p < 1 x 10^-4^). This correlation has been associated with an effect of recombination in driving GC through GC biased gene conversion but the relatively low correlation coefficients could indicate that this force is not strong in termites. By contrast, the correlation between GC content and recombination rate is much stronger in honey bee (R^2^ = 0.506), which is likely caused by high recombination rates and intense GC-biased gene conversion (Wallberg et al. 2015).

There is a weak but significant correlation between overall repeat element density and recombination rate in *C. secundus* (Spearman’s *ρ* = = 0.02, p < 1 x 10^-4^) but not for *M. bellicosus*. There is a weak but significant correlation between simple repeats and recombination in both genomes (Spearman’s *ρ* = 0.051, p < 2.22 x 10^-16^ for *M. bellicosus* and Spearman’s *ρ* = 0.055, p < 2.22 x 10^-16^ for *C. secundus*). SINE and LINE elements are also only weakly associated with recombination rate in both genomes (SINEs: Spearman’s *ρ* = -0.012, not significant for *M. bellicosus* and Spearman’s *ρ* = 0.02, p < 1 x 10^-4^ for *C. secundus*; LINEs: Spearman’s *ρ* = -0.02, p < 1 x 10^-4^ for *M. bellicosus* and Spearman’s *ρ* = 0.002 not significant for *C. secundus*). There is a significant negative correlation between gene density and recombination for the *M. bellicosus* genome (Spearman’s *ρ* = -0.185, p < 2.22 x 10^-16^), but no significant correlation for *C. secundus*. We also tested whether hotspots differ from the background with respect to GC content. The GC content does not differ substantially between hotspots and the rest of the genome in either species (means of 40.1% and 41.0% for *M. bellicosus* and 40.2% and 40.3% for *C. secundus*).

We found a significant correlation between nucleotide diversity (π) and recombination in both species (Supplemental Figure S2), with Spearman’s *ρ* = 0.48, p < 1 x 10^-4^ for *M. bellicosus* and Spearman’s *ρ* = 0.31, p < 1 x 10^-4^ for *C. secundus*. This correlation is found across a wide range of eukaryotic taxa and is generally accepted to be caused by the interaction between recombination and linked selection (Nachman 2001; Begun and Aquadro 1992).

### Recombination and gene expression patterns correlate with CpG_O/E_

Analysis of the distribution of CpG_O/E_ in both termite species showed that it is bimodally distributed, with substantially lower values in introns and exons compared to flanking noncoding regions (Supplemental Figure S3). This indicates that germline DNA methylation is biased towards gene bodies in the two termite genomes, a pattern that is also found in honey bees and other insect genomes with functional DNA methylation (Harrison et al. 2018; Elango et al. 2009; Sarda et al. 2012). We found that exons have lower CpG_O/E_ compared to introns, which are both lower than flanking regions (Figure 5, top row; all comparisons p < 0.001 after Bonferroni correction) consistent with higher levels of germline methylation in exons (Supplemental Table S6). We next analysed how recombination rate varies among genomic features. We found that *ρ*/kbp is also significantly reduced in genes compared to flanking regions for *C. secundus,* with exons also showing reduced *ρ*/kbp compared to introns (Figure 5, bottom row, all p-values <0.005 after Bonferroni correction). The same trends are observed in *M. bellicosus* although they are not significant. The correlation between CpG_O/E_ and recombination rate among genes is higher than for the genome overall (Spearman’s *ρ* = 0.315, p < 2.22 x 10^-16^ for *M. bellicosus*; Spearman’s *ρ* = 0.196, p < 2.22 x 10^-16^ for *C. secundus*).

**Figure 5.**
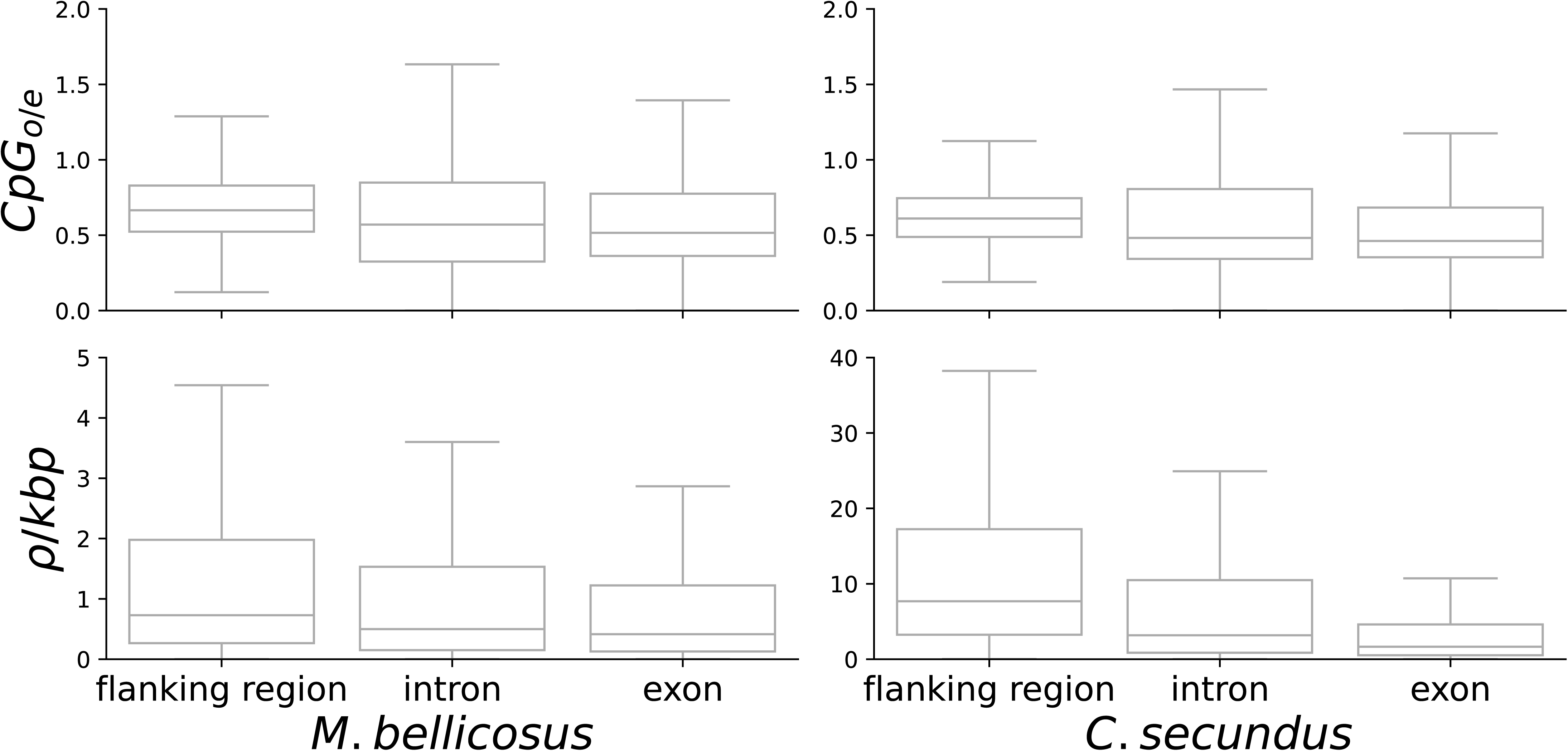
Boxplots showing variation in levels of CpG_O/E_ and recombination rate (*ρ*/kbp; rho) in genic and flanking regions in the *M. bellicosus* and *C. secundus* genomes. The differences between all categories are significant.

We next investigated whether patterns of gene expression between castes and sexes (hereafter, for short ‘caste-biased gene expression’) were correlated with CpG_O/E_ and recombination rate using gene expression data from two studies (Lin et al. 2021; Elsner et al. 2018). It has been proposed that it is evolutionarily advantageous for genes with worker-biased expression to be located in regions with elevated recombination (Kent et al. 2012). Comparisons of CpG_O/E_ and recombination rate for the original gene expression categories reported by (Elsner et al. 2018) for *M. bellicosus* are shown in Supplemental Figure S4. We also reclassified the genes into the following expression categories: queen-biased, king-biased, worked-biased, male-biased, female-biased, reproduction-biased, and differentially expressed (see methods for details). For *C. secundus*, only worker-biased and queen-biased genes were identified (Lin et al. 2021). We find that mean CpG_O/E_ in gene bodies shows significant variation between gene expression categories in both *M. bellicosus* and *C. secundus* (Figure 6). However, this variation is not consistent between species. In *M. bellicosus*, queen-biased and female-biased genes have low CpG_O/E_ and king-biased and male-biased genes have high CpG_O/E_. In *C. secundus*, queen-biased genes have higher than average CpG_O/E_.

**Figure 6.**
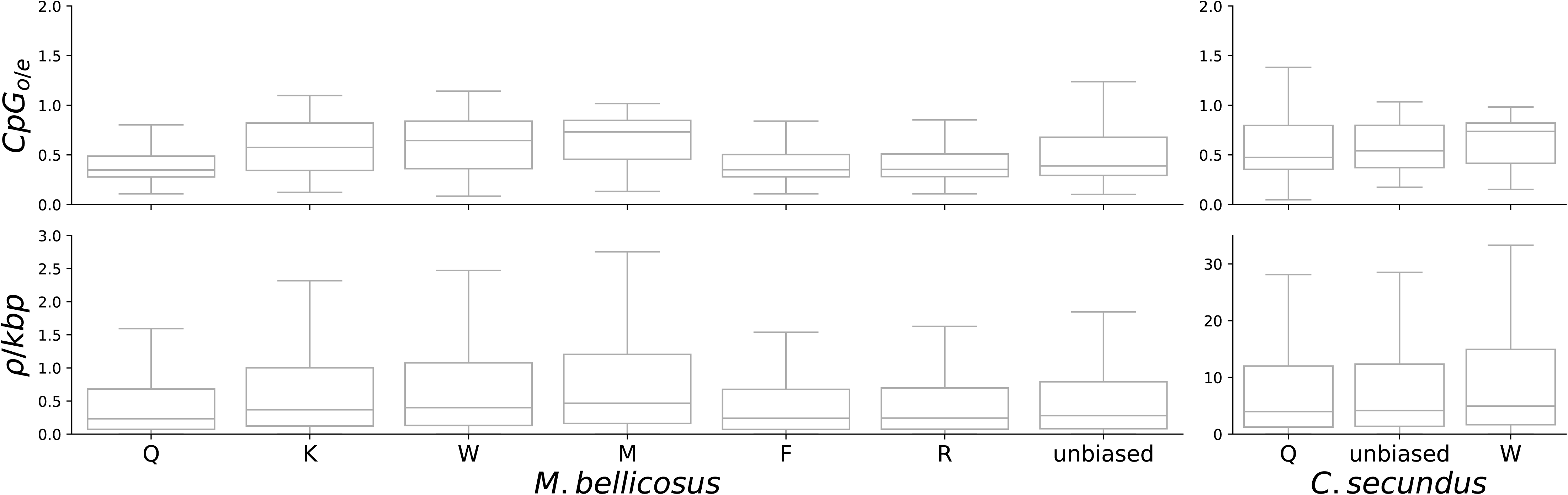
Boxplots showing variation in levels of CpG_O/E_ and recombination rate (*ρ*/kbp) in the coding region of genes in the *M. bellicosus* and *C. secundus* genomes classified according to their patterns of gene expression (Q = queen-biased, K = king-biased, W = worker-biased, F = female-biased, R = reproductive-biased).

The differences in CpG_O/E_ between caste-biased gene expression categories are mirrored by differences in *ρ*/kbp (Figure 6). Gene expression categories with elevated CpG_O/E_ consistently show elevated *ρ*/kbp, and this pattern is found in both species. Studies in the honey bee have found that worker-biased genes have lower CpG and higher recombination rates (Kent et al. 2012; Wallberg et al. 2015). In *M. bellicosus,* we find that worker-biased genes have slightly elevated CpG_O/E_ and *ρ*/kbp. However, in *C. secundus*, these values are reduced in worker-biased genes, whereas queen-biased genes have a slightly elevated CpG_O/E_ and *ρ*/kbp in this species.

We investigated the relative roles of CpG_O/E_ and gene expression in determining *ρ* in genes using linear models, with CpG_O/E,_ *ρ* in flanking region and expression categories as explanatory variables. In both *M. bellicosus* and *C. secundus*, we found that CpG_O/E_ was the strongest predictor of recombination rate variation among genes (Supplemental Tables S7 and S8) in line with the analyses above. We found that caste-biased expression had no significant impact on the recombination rate. CpG_O/E_ is a better predictor of *ρ*/kbp in genes than *ρ*/kbp in flanking region and variation in *ρ*/kbp among expression categories mainly reflects differences in CpG_O/E_.

## Discussion

The main findings we present here are 1) that two distantly related termite species with varying social complexity both have relatively low genomic average rates of meiotic recombination, 2) both genomes possess recombination hotspots and likely contain full-length copies of *PRDM9* and, 3) there are reduced levels of recombination in genomic regions depleted in CpG sites. Our findings contrast with the extremely elevated recombination rates observed in eusocial Hymenoptera, which are likely to be generated by a feature that is specific to these taxa. The finding of recombination hotspots is one of the first in insects. This could reflect a conserved function of PRDM9 in initiating recombination events in both vertebrates and invertebrates. Our results also support a role for germline methylation in suppressing recombination.

### Low recombination rates in termites

Data from a wide range of taxa indicate that at least one crossover per chromosome tetrad is essential for correct chromosomal segregation during meiosis (Fernandes et al. 2018). The chromosome numbers of *M. bellicosus* and *C. secundus* are 2n = 42 and 2n = 40, respectively (Jankásek et al. 2021). Considering their assembly lengths, this indicates that a minimum average genomic recombination rate of 0.92 cM/Mb and 1.01 cM/Mb, respectively, would be necessary for accurate meiosis to occur in the two termite species. Our estimates of recombination rate per meiosis based on analysis of LD depend on several factors, including the algorithm to estimate *ρ* and the estimate of mutation rate. As no estimates of the mutation rate in termites were available, we considered a range of estimates from other insects. These estimates vary by more than an order of magnitude between different insect species (Lynch et al. 2023) and higher estimates of the mutation rate give higher estimates of the recombination rate. Our estimates based on LDhat are all less than 1 cM/Mb for both species, even when considering the highest estimate of mutation rate. Using estimates from LDhelmet and the average of experimentally determined mutation rate estimates in insects gives estimates of recombination rate of 1-2 cM/Mb. The species most closely related to termites for which a mutation rate estimate was available was *H. coronatus (Huang et al. 2023)*. Using this estimate and *ρ* from LDhelmet, the recombination rate per meiosis was estimated to 0.57 cM/Mb for *M. bellicosus* and 0.69 cM/Mb for *C. secundus*, i.e. both slightly lower than expected based on the requirement of one crossover per chromosome. Taken together, these results indicate that recombination rates in these two termite species are substantially lower than the elevated values found in eusocial Hymenoptera (6-25 cM/Mb) (Wilfert et al. 2007).

It should be noted that our estimates of recombination rate are sex-averaged rates. It is possible that recombination is absent in one of the termite sexes, which would make a lower sex-averaged recombination rate more feasible. In addition, sex-linked contigs have not been identified in the termite genome assemblies utilized here. *M. bellicosus* and *C. secundus* have X1X2/Y1Y2 and X/Y sex-determining systems respectively (Jankásek et al. 2021). The inclusion of sex chromosomes in the analysis leads to errors in determining the genome-wide recombination rate per meiosis because they have lower *N*_*E*_ than the rest of the genome. In addition, sex chromosomes are more prone to assembly and read mapping errors, due to reads aligning with the incorrect copy of the chromosome. However, considering that sex chromosomes only comprise a minor portion of the genome, this is unlikely to substantially alter our results. It has also been observed that some chromosomes form ring or chain structures during meiosis in certain termite species (Bergamaschi et al. 2007), which could plausibly relax the requirement for one crossover per chromosome per meiosis.

### Eusociality is not a universal driver of high rates of meiotic recombination

Elevated genomic rates of recombination are characteristic of all eusocial Hymenoptera so far investigated. Several hypotheses have been advanced to explain this phenomenon, which are all based on an evolutionary advantage of recombination in eusocial species (Wilfert et al. 2007). For instance, it has been suggested that elevated recombination could increase genetic diversity in a colony. This could promote diversity in the workforce, enabling more efficient task specialisation among workers (Kent et al. 2012). Alternatively, it could render the colony less susceptible to invasion by parasites and pathogens (Fischer and Schmid-Hempel 2005). Elevated recombination might also favour rapid evolution of caste-specifically expressed genes (Kent and Zayed 2013). In addition, increased recombination could lead to reduced variance in relatedness, which could prevent kin conflict in colonies (Sherman 1979; Templeton 1979).

All these hypotheses predict that recombination rate should be elevated in all eusocial insects. Here, however, in the first report of recombination rate in social insects outside of Hymenoptera, we find overall low levels of recombination. Both termite species studied here are eusocial, and in particular *M. bellicosus* has extremely large and complex colonies characteristic of eusociality (Korb and Thorne 2017). The low recombination rate inferred in the two termite genomes prompts re-evaluation of hypotheses connecting eusociality with high recombination rates. Although both termites and social Hymenoptera share many traits, they differ in others that reflect their different ancestries.

The chromosome numbers of both termite species (*M. bellicosus*, 2n = 42; *C. secundus*, 2n = 40) are typical for termites, although variation exists between taxa (Jankásek et al. 2021). These values do not differ from those observed in Hymenoptera, which have a similar range (e.g. *A. mellifera*, 2n = 32) and do not vary between solitary and eusocial taxa (Cardoso et al. 2018; Cunha et al. 2021; Ross et al. 2015). This suggests that differences in chromosome number are unlikely to be relevant in explaining the differences in recombination rates among these taxa.

One proposed advantage of sex and recombination is that it is associated with sexual selection and a higher intensity of selection on males. This could be advantageous because differential male mating success can reduce mutational load in sexual populations by causing deleterious mutations to segregate at a lower equilibrium frequency on average (Siller 2001; Agrawal 2001). Recombination is essential for effective purging of deleterious alleles as it unlinks deleterious variants from nearby beneficial ones thus preventing selection interference (Hartfield and Keightley 2012). Selection has been found to be stronger in males than females across animal species (Janicke et al. 2016; Winkler et al. 2021). Species with a male-biased sex ratio are expected to experience particularly intense selection on males. It is therefore possible that recombination rate could be particularly elevated in species in which males face intense competition as this increases the evolutionary advantage of recombination.

Another proposed advantage of sex and recombination in general is that it is associated with the intensity of sperm competition. Maynard Smith (1976) presented a model that explains how sib-competition can produce an immediate evolutionary advantage to sex and recombination, which occurs when several offspring of a single female compete within a patch. This model was extended by Manning and Chamberlain (1997) who argued that it is analogous to intra-ejaculate competition among sperm, in which recombination increases the variance of fitness. Under these conditions, if inter-ejaculate competition also exists, it is predicted that there will be selection for increased recombination rate. This model predicts a significant correlation between recombination rate and gamete redundancy (excess sperm production), which is observed in mammals (Manning and Chamberlain 1997).

Many social Hymenoptera produce highly male-biased reproductive offspring and thousands of haploid male drones compete to fertilise a small number of female queens (Winston 1991; Koeniger et al. 2011; Baer 2005). This situation is similar to sperm competition as haploid males can be considered analogous to gametes. However, as many more genes are expressed in adult haploid males compared to gametes there are more targets of selection. Selection against deleterious alleles is particularly effective in haploid males because recessive deleterious alleles are exposed (Hedrick and Parker 1997). It is therefore possible that the male-biased sex ratio observed in many eusocial Hymenoptera could favour elevated recombination rates in the species so far studied, due to elevated competition between males.

Male-biased sex ratios are particularly strong in the genus *Apis*, which is also the genus in which the most extreme recombination rates have been inferred (Beye et al. 2006; Shi et al. 2013; Liu et al. 2015; Wallberg et al. 2015; Rueppell et al. 2016; Kawakami et al. 2019). Male-biased sex ratios are also observed in other social bees and wasps. Both male and female biased operational sex ratios have been observed in ants, but are difficult to measure because workers invest more resources in female offspring and may manipulate sex ratios. In non-social Hymenoptera, females lay roughly equal numbers of haploid and diploid eggs (Trivers and Hare 1976). The sex ratio of reproductive males and females is also relatively even in most termite species (Roisin 2001).

Further research is needed to determine the cause of variation in recombination rates among social and non-social insects. For example, simulations could be used to determine the effect of selection on haploid males on recombination rates, considering different sex ratios. In addition, it is necessary to estimate recombination rate in a wider range of social and non-social species to determine whether the strength of selection on males is a key factor in determining these rates.

### Full length PRDM9 gene and recombination hotspots in termite genomes

Our results are also indicative of the presence of recombination hotspots in the two termite genomes. The fine-scale distribution of recombination rate in both termite genomes shows that 50% of the recombination happens in less than 0.5% of the genome, which is even more extreme than for humans where this corresponds to ∼6% of the genome (Myers et al. 2005). This contrasts to the honey bee and solitary bees where fine-scale maps are available and recombination is relatively uniformly distributed in the genome (Wallberg et al. 2015; Jones et al. 2019).

Two main mechanisms have been shown to lead to recombination hotspots. The first, which is best understood in human and mouse, is governed by the PRDM9 protein. A zinc-finger domain recognises a specific DNA sequence motif and then performs a histone modification in the vicinity, which marks the sequence for a DNA double-stranded break that is repaired by recombination during meiosis (Baudat et al. 2010; Myers et al. 2010; Parvanov et al. 2010). In species with this mechanism, both the PRDM9 zinc-finger domain and the sequences present in hotspots are fast evolving, which results in a rapid turnover of hotspot locations (Myers et al. 2010). Full-length PRDM9 orthologues are present among a range of highly-diverged vertebrates including many fish, reptiles and mammals, but many full or partial losses of the gene have also been inferred. In a study of 225 vertebrate genomes, a minimum of six partial and three complete losses were inferred (Baker et al. 2017). However, it is unclear whether PRMD9-hotspots are common in invertebrates.

A second mechanism is found in vertebrates that lack PRDM9, in which recombination hotspots are localised to regions of open chromatin such as CpG islands and promoters. This is found in dogs and in most or all birds, in which PRDM9 has been lost, and results in hotspots with stable rather than rapidly evolving locations (Axelsson et al. 2012; Singhal et al. 2015). A similar pattern is also observed in the yeast *Saccharomyces cerevisiae* in which recombination preferentially occurs in promoter regions (Wu and Lichten 1994). PRDM9-independent hotspots can be identified in CpG islands in some vertebrate genomes even when PRDM9 is present (Joseph et al. 2024). In invertebrates such as *Drosophila* and *Caenorhabditis elegans*, recombination hotspots appear to be absent (Coop and Przeworski 2007; Chan et al. 2012; Smukowski Heil et al. 2015). In insects, fine-scale maps of social and solitary bees have not revealed recombination hotspots (Wallberg et al. 2015; Jones et al. 2019), but there is evidence for hotspots in a butterfly genome (Torres et al. 2023).

We find evidence that recombination is directed towards hotspots in both termite species. We also identify a full-length *PRDM9* gene in both species, containing all of the domains needed to initiate recombination. However, we are unable to demonstrate that PRMD9 is responsible for the presence of recombination hotspots in the two genomes. We used a motif prediction algorithm to predict the zinc-finger binding motif in *M. bellicosus* and *C. secundus*, but we did not observe elevated recombination rate around instances of the motif in either of the genomes. Furthermore, we did not identify any common sequence motifs that were strongly enriched in recombination hotspots in either of the termite genomes. A plausible interpretation of these observations is that the PRDM9 protein indeed initiates recombination in hotspots in both species, but that positive selection leads to frequent shifts in the target motifs, which results in a lack of a strong association between a specific motif and LD-based hotspots (Myers et al. 2010). It is also possible that the sequences of the zinc-finger motifs are not correctly represented in the genome assemblies, which could result from problems with assembly around minisatellites. However, an unknown mechanism could also be responsible for hotspots in termites.

### Evidence that germline DNA methylation suppresses recombination

In insects with a functioning DNA methyltransferase machinery, the CpG_O/E_ statistic shows substantial variation along the genome, which mainly reflects variation in levels of germline DNA methylation (Elango et al. 2009). In contrast to vertebrate genomes, methylation is strongly skewed towards gene bodies in the genomes of insects (Glastad et al. 2011). In the two termite genomes studied here, we find lowest values of CpG_O/E_ in exons, whereas noncoding flanking regions are biased towards higher values, indicative of lower levels of DNA methylation (Figure 5, Supplemental Figure S3). Exons, introns and noncoding flanking regions all display a predominantly bimodal distribution of CpG_O/E_, likely reflective of the presence of methylated and unmethylated regions. We find that recombination rate is reduced in gene bodies compared to flanking regions in one of the two termite genomes, but the overall genomic correlation between CpG_O/E_ and recombination rate, although significant, is weak. This could reflect the fact that the majority of the genomes are not methylated, so that much of the variation in CpG_O/E_ on the genome scale is not strongly influenced by methylation. Our inference of reduced recombination rates in exons could also be influenced by lower *N*_*E*_ in these regions due to the effects of linked selection, which could result in more extensive LD (Charlesworth 2009). However, the observation that differences in CpG_O/E_ among genes are associated with recombination rate indicates that linked selection is unlikely to be a major factor driving the observed differences in recombination rate and that germline DNA methylation is more important.

We identified CpG_O/E_ as the factor with the strongest influence on recombination rate variation among genes. There were no significant effects of differences in gene expression patterns across castes, a result that was also observed in the honey bee genome (Wallberg et al. 2015). This indicates that recombination rate variation is not modulated by differences in gene expression in these genomes, which might be expected if natural selection favours increased recombination rates in certain genes. For example, it has been proposed that natural selection could favour elevated recombination rates in genes with roles in worker behaviour or in immune function due to their important roles in colony function (Wilfert et al. 2007; Kent et al. 2012). However, the results presented here suggest that differences in recombination rate between genes with caste-biased patterns of gene expression are mainly due to underlying differences in methylation levels among genes.

A direct effect of germline DNA methylation in suppressing recombination could influence variation in recombination landscapes among vertebrates and invertebrates. Vertebrate genomes are usually highly methylated with the exception of CpG islands. In vertebrate genomes that lack a functional *PRDM9* gene, recombination events tend to localise to CpG islands (for example in dogs and birds) (Singhal et al. 2015; Axelsson et al. 2012). The effect of DNA methylation on recombination rate variation in invertebrates is less clear, but the results presented here also support a role of germline methylation in suppressing recombination.

## Materials and Methods

### Sample collection

*M. bellicosus* samples were collected from colonies in Kakpin, next to the Comoé National Park in Côte d’Ivoire (coordinates 8°3911N 3°4611W) (Elsner et al. 2018). *C. secundus* colonies were collected from dead *Ceriops tagal*mangrove trees near Palmerston-Channel Island (Darwin Harbor, Northern Territory, Australia; 12°3011S 131°0011E). Colonies were then kept in *Pinus radiata* wood blocks in climate rooms in Germany, providing 28 °C, 70% relative humidity, and a 12-h day/night cycle (Lin et al. 2021). For each species, each of the 10 collected samples came from a different colony.

### Population sequencing

The termites were cut in two, approximately between head and thorax. DNA was extracted from both parts using the QIAGEN Blood and Tissue Kit following the standard protocol, including treatment with 4 µL RNase and 25 µL Proteinase K. The DNA extraction product was cleaned and concentrated with ZymoResearch Genomic DNA Clean & Concentrator kit. For most samples, the RNase treatment was repeated prior to cleaning and concentrating. One DNA sample from each individual was chosen, based on the DNA amount and quality, to be sequenced. Sequencing libraries were produced using the Illumina DNA PCR-free library preparation kit according to the manufacturers protocol, using an input DNA quantity of 200 ng per sample. This protocol yields insert sizes of approximately 350 bp. We performed Illumina short-read sequencing on the libraries using NovaSeq 6000 S4 flow cell with 2 x 150 bp paired end reads.

### Read mapping and variant calling

The sequence data from the two termite species were mapped to the reference genomes for *M. bellicosus* (Qiu et al. 2023) and *C. secundus* (Csec_1.0) (Harrison et al. 2018) respectively, using Burrows-Wheeler alignment tool BWA version 0.7.17 (Li & Durbin 2009) with the BWA-MEM algorithm. Sorting and indexing of the BAM-files was done with the SAMtools package version 1.17 (Li et al. 2009), followed by adding read groups and marking duplicate reads with Picard toolkit version 1.118 (http://broadinstitute.github.io/picard/). After mapping, we excluded all scaffolds < 1 kbp from the *C. secundus* genome, which in total correspond to 2% of the genome size, as they are potentially of lower quality and not informative for analysis of linkage disequilibrium or recombination. For *M. bellicosus* there are no scaffolds shorter than 1 kbp, but we excluded the following five scaffolds, which together correspond to 3.78% of the genome length, due to unexpectedly high heterozygosity: 93, 62, 41, 35 and 32.

One of the *C. secundus* individuals did not map well to the reference genome (71% mapped reads, unequal distribution between forward and reverse strand and an average mapping quality of 35 while all the other samples had an average mapping quality >45). This individual was excluded from all further analyses and hence the total number of termite individuals analysed in this study is 19, including 10 *M. bellicosus* and 9 *C. secundus*.

Variant calling was done using the tools HaplotypeCaller, GenomicsDBImport and GenotypeGVCFs from GATK version 4.3.0.0 as described in their best practices workflow (https://gatk.broadinstitute.org/hc/en-us/articles/360035535932-Germline-short-variant-discovery-SNPs-Indels-, last accessed 2023-07-11). The variants were filtered in multiple steps, first removing extended regions of low mapping quality or deviating read depth as those regions are likely to be unreliable. This was done based on statistics from the SAMtools mpileup and depth tools, respectively, removing 100 kbp windows with mean mapping quality below 70 or mean read depth more than two standard deviations from the genome-wide mean. Indels were removed using VCFtools version 0.1.16 (Danecek et al. 2011) before GATK VariantFiltration was run with the following limits, which are all equal to or stricter than the ones recommended (https://gatk.broadinstitute.org/hc/en-us/articles/360035890471-Hard-filtering-germline-short-variants, last accessed 2023-07-11) and adapted to the observed distributions of the corresponding statistics: QD<2; FS>50; MQ<40; ReadPosRankSum< -4; ReadPosRankSum>4; SOR>3; ExcessHet>5 for both species as well as MQRankSum with limits -5 and 5 for *C. secundus* and limits -6 and 4 for *M. bellicosus*. This was followed by additional filtering with VCFtools to select only bi-allelic sites with a minimum quality score of 30 and maximum of 40% missing genotypes. The SNPs with mean read depth in the lowest or highest 2.5% of the depth distribution were also filtered out, which translates to a depth below 10 or above 34 for *C. secundus* and below 5 or above 40 for *M. bellicosus.* For the recombination rate calculations, the SNPs were also filtered for a minor allele count >= 2, as rare variants appearing on only one chromosome have a relatively high false positive rate and are not informative regarding LD. The average proportion of missing genotypes per individual was 4.5 x 10^-5^ for *C. secundus* and 1.2 x 10^-4^ for *M. bellicosus*. After variant calling and filtering, the data was phased and imputed using Beagle version 5.1 (Browning and Browning 2007; Browning et al. 2018).

### Estimation of LD decay

The decay of linkage disequilibrium, *r*^2^, with increasing physical distance was estimated with the software PopLDDecay version 3.42 (Zhang et al. 2019) over a maximum distance of 10 kbp. For this analysis, sequence data from *A. mellifera scutellata* was included for comparison (Wallberg et al. 2017). The *A. m. scutellata* sequences were mapped to the Amel_HAv3.1 reference genome (Wallberg et al. 2019) and processed and filtered in the same way as described above for the termite data (GATK VariantFiltration with the limits QD<2, FS>25, MQ<40, MQRankSum<-4, MQRankSum>4, ReadPosRankSum<-4, ReadPosRankSum>4, SOR>3, ExcessHet>5 and VCFtools to select bi-allelic sites with mean read depth between 5-13, quality score >= 30, minor allele count >=2 and <=40% missing genotypes). For *M. bellicosus*, the reference genome contains some gaps of unknown size and in order to avoid estimating *r*^2^ across those gaps, scaffold breaks were introduced at the corresponding positions.

### Estimation of population recombination rate

We estimated the population-scaled recombination rate, *ρ,* using two methods: LDhat (Auton and McVean 2007) and LDhelmet (Chan et al. 2012), which are both based on a Bayesian reversible-jump Markov Chain Monte Carlo algorithm (rjMCMC).

For LDhat (https://github.com/auton1/Ldhat, downloaded 2023-04-19), the input files were generated from the filtered vcf-file with VCFtools and the --ldhat option. Then the LDhat function complete was run in order to create a lookup table, with the estimated value of *θ*_*W*_ (see below), a maximum *ρ* value of 100 and 101 grid points, as recommended. This was followed by the main function, interval, to perform the rjMCMC with 10 million iterations, sampling every 5000th iteration and a block penalty of 1 as this has been used previously with LDhat in similar studies (Wallberg et al. 2015; Jones et al. 2019). In order to summarise the output from interval, the stat function was used, discarding the first 20 samples as burn-in.

The recombination rate was also estimated using the software LDhelmet version 1.10 (Chan et al. 2012). In order to prepare the input files for LDhelmet, the filtered vcf-files were converted into multiple-sequence fasta-files using the vcf2fasta function from vcflib version 2017-04-04 (Garrison et al. 2022) and to the related file formats .snps and .pos with the - - ldhelmet option from VCFtools (version 0.1.16). The first step of the LDhelmet workflow is to create haplotype configuration files with the find_confs tool, followed by generation of likelihood lookup tables with the tool table_gen and computation of Padé coefficients with the tool pade. These tools were run with the recommended parameters, meaning that find_confs was run with a window size of 50 SNPs, table_gen was run with a grid of *ρ-*values ranging from 0 to 100 with increments of 0.1 up to 10 and increments of 1 for the remaining grid and pade was run to generate 11 Padé coefficients. Scaffolds shorter than or equal to the window size of 50 SNPs were removed, corresponding to 1% of the sequence length in *C. secundus* and even less for *M. bellicosus*. The tool rjmcmc which contains the main algorithm was run with a burn-in of 100,000 followed by 1,000,000 iterations with a block penalty of 50 as recommended (Chan et al. 2012). For each species, the rjmcmc tool was run three times with different seeds and the convergence between the replicates was evaluated in terms of Spearman’s rank correlation coefficient, which was > 0.95 with p < 2.2 x 10^-16^ for all pairwise comparisons of both species. The binary output from rjmcmc was converted to text, extracting the mean *ρ-*value between each pair of SNPs, using the tool post_to_text.

The outputs from LDhat and LDhelmet give the value of the parameter *ρ* between each pair of consecutive SNPs. Those values were converted to equally sized windows across each scaffold, as well as the coordinates of individual genomic features, by taking a weighted average of values from overlapping intervals. The *M. bellicosus* scaffolds contain a few gaps of unknown size and the values between SNPs across such gaps were removed.

The parameter *ρ* is proportional to the recombination rate *r* and the effective population size *N*_*E*_, which in turn can be estimated based on the mutation rate *μ* and the number of segregating sites, *K*. In order to convert *ρ* to *r*, the following equations were used, where *n* is the number of haploid sequences. The number of segregating sites (*K*) was calculated before the filter on minor allele count >=2 was applied, as this filter likely removes many true SNPs as well, which could affect the estimates.

*θ*_*W*_ was estimated for each sample using the equation below from (Watterson 1975):

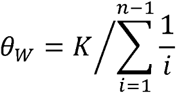

We used the value of *θ*_*W*_ to estimate *N*_*E*_ using:

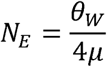

We estimated *N*_*E*_ using a range of mutation rates estimated from other insect species (Keightley et al. 2009, 2014, 2015; Yang et al. 2015; Liu et al. 2017; Lynch et al. 2023). We then estimated the recombination rate using the following equation, assuming a range of values of *N*_*E*_.

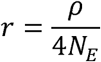

### Cumulative plot of rho *ρ*/kbp

A cumulative plot of the proportion of recombination events versus the proportion of the physical distance along the genome was constructed based on estimates of *ρ*/bp for the two termite species and previous estimates for *A. mellifera* (Wallberg et al. 2015; PRJNA236426). The *ρ*/bp estimates between markers were placed in decreasing order before multiplication with the respective physical distances between the markers. Then cumulative sums were calculated for the *ρ*-values as well as the corresponding physical distances and divided by the total genome-wide sums.

### k-mer enrichment in recombination hotspots

In order to identify *k*-mers associated with elevated recombination rate, we defined recombination hotspots as 2 kbp segments with at least five-fold elevated recombination rate compared to the local background, as defined by 50 kbp flanking up and downstream of the segment, resulting in 39837 regions for *M. bellicosus and* 40894 regions for *C. secundus*.

Then, we used these hotspots to search for motifs associated with elevated recombination rate by comparing against an equivalent number of randomly chosen 2 kbp regions from the rest of the genome. This was done using STREME 5.5.4 (Bailey 2021) using a minimum and maximum size of *k*-mer of 8 and 20, ( --minw 8 ; --maxw 20) and otherwise default options. From all motifs with significant enrichment after adjusting for multiple testing, we then considered only motifs present in more than 5% of all hotspots and an at least two-fold enrichment compared to the background as credible candidates.

### Identifying PRDM9 orthologues in M. bellicosus and C. secundus

A full-length copy of PRDM9 is present in the *C. secundus* genome annotation, but not in *M. bellicosus*. Using OrthoFinder (Emms and Kelly 2019), and the genome annotation files for both species, we identified a single orthologue to the *C. secundus* PRDM9 in *M. bellicosus*. We analysed this region using the NCBI conserved domain search (Wang et al. 2023). We also searched the *M. bellicosus* assembly for the PR/SET domains derived from *C. secundus* and *Zootermopsis nevadensis* using BLAST (Camacho et al. 2009) and subsequently for all other PRDM9 domains (KRAB, SSXRD, ZF-Casette) to identify hits that contain all other PRDM9 domains in the vicinity in the correct order. We used GeneWise (Madeira et al. 2024) with default options to perform gapped alignment of a the PRDM9 protein sequence from *Zoothermopsis nevadensis* against the target region to predict exon-intron borders.

### Predicting binding sites from zinc-finger-motifs

Using the ZF-motifs from *C.secundus,* we obtained letter probability matrices for predicted binding motifs using a webserver (http://zf.princeton.edu/index.php; (Persikov and Singh 2014) which were used to search for binding sites across the genome using FIMO (Grant et al. 2011) with default options. Subsequently, the mean rho for the surrounding 1 kbp bin was used to test for a difference in mean rho between bins associated with a suspected binding site, and a similar number of randomly chosen bins without association. For significance testing, we repeated this draw 10.000 times to compare the associated bins against the 95% confidence interval of the resulting distribution.

### Analysis of genomic correlates of recombination

We divided the genome into 10 kbp windows in which average *ρ* and other genomic features were estimated. Windows shorter than 5 kbp (half of the specified window size), which appear in short scaffolds or at the ends at longer scaffolds, were excluded from the analyses (excluding less than 1% of the sequence for each species). Correlation coefficients were estimated with Spearman’s *ρ*, with significance estimated by permutation tests with 10,000 iterations.

In order to determine the repeat content across the *M. bellicosus* and *C. secundus* genome assemblies, we first generated custom repeat libraries for each species with RepeatModeler (Flynn et al. 2020) using default options. This was then combined with Repbase 29 (Bao et al. 2015) and Dfam 3.8 (Storer et al. 2021) and used to run RepeatMasker with default options on each genome. We estimated the proportion of sequence in each repeat class in 10 kbp windows and used Spearman’s rank correlation to correlate this with estimates of recombination rate.

### Per gene recombination rate and CpG_O/E_ ratio

The observed/expected frequency of CpG sites (CpG_O/E_) and mean *ρ* were calculated on a per-gene basis, as well as for exons and introns, and 50 kbp flanking regions 10 kbp upstream and downstream of the gene, akin to Wallberg et al (2015). For recombination rate, estimates of *ρ* between markers were used, utilising a weighted mean when the element spanned multiple markers. For visualisation, extreme CpG_O/E_ values due to low GC content were set to a maximum of 4.

We tested for significant difference in mean between *ρ* and CpG_O/E_ in flanking regions, exons and introns using paired *t*-tests corrected for multiple testing using a Bonferroni threshold.

### Differential expression data

We classified genes in both species according to their caste-biased patterns of expression from two studies (Elsner et al. 2018; Lin et al. 2021). Elsner et al. (2018) analysed four castes of *M. bellicosus*, with two age classes of each: major and minor workers, queens and kings. They analysed differences in gene expression among these classes using RNA-seq data. We reclassified the differential expression data by grouping the direct caste-vs-caste comparisons as "queen-biased", “king-biased”, “worker-biased”, “reproduction-biased”, "male-biased" or "female-biased" if they were differentially upregulated for this caste (or group) in comparison to other castes (or groups). Since Elser *et al*. (2018) initially mapped the *M. bellicosus* reads against the *Macrotermes natalensis* genome, We used OrthoFinder (Emms and Kelly 2019) to identify the *M. bellicosus* orthologues corresponding to the differentially expressed genes. Only single-copy genes were considered for this analysis.

Lin et al. (2021), performed RNA-seq on samples of workers and queens. Here, we used the data from untreated queens and workers. Differentially expressed *C. secundus* genes were classified as “worker-biased” or “queen-biased” if they were differentially expressed with a bias to the respective caste, or unbiased if they were not, and subsequently compared between classes. The lists of genes we identified in the caste-biased categories in both species are provided as Supplemental Tables S9 and S10.

### Correlates of recombination rate variation among genes

In order to assess the relative importance of CpG_O/E_ and differential expression patterns on recombination rate in genes (genic *ρ*/kbp), two ordinary least squares models were built using statsmodels (Seabold and Perktold 2010) for each termite species. Genic CpG_O/E_, differential expression categories and flanking region *ρ*/kbp were used as exogenous variables. In the first model, we included flanking region *ρ*/kbp and CpG_O/E_. In the second model, we included all available biased expression categories for each species.

## Data access

The Illumina whole-genome sequencing data generated in this study have been submitted to the NCBI BioProject database (https://www.ncbi.nlm.nih.gov/bioproject/) under accession number PRJNA1021607. Custom scripts are available on GitHub (https://github.com/troe27/termite-recombination-rate) and as Supplemental Code.

## Competing interest statement

The authors declare no competing interests.

## Acknowledgements

We acknowledge grant 2018-03896 from The Swedish Research Council to MTW and from the Deutsche Forschungsgemeinschaft (DFG) KO-1895/26-1: 417980976 to JK. We acknowledge support from the National Genomics Infrastructure in Stockholm funded by Science for Life Laboratory, the Knut and Alice Wallenberg Foundation and the Swedish Research Council. The computations were enabled by resources in project NAISS 2023/23-402 provided by the National Academic Infrastructure for Supercomputing in Sweden (NAISS) at UPPMAX, funded by the Swedish Research Council through grant agreement no. 2022-06725. We thank Göran Arnqvist for helpful discussions on the evolution of sex and recombination and Marie Raynaud for assistance running LDhelmet. The Parks and Wildlife Commission, Northern Territory, and the Department of the Environment, Water, Heritage and the Arts, Australia, gave permission to collect (Permit number 59044) and export (PWS2016-001559) the *C. secundus* termites. The Office Ivoirien des Parcs et Réserves (OIPR) provided sampling and export permits for the *M. bellicosus* samples. The study was conducted according to the Nagoya protocol. MTW designed and led the research; TE and TR conducted the analysis; MTW, TE, TR and JK wrote the paper; DE and JK contributed data and samples; AO, YL, and TL performed lab work.

## Notes

### Competing Interest Statement

The authors have declared no competing interest.

### Summary of Updates

small typos corrected, methods section improved

