## supplemental figure for "Unexpectedly low recombination rates and presence of hotspots in termite genomes"

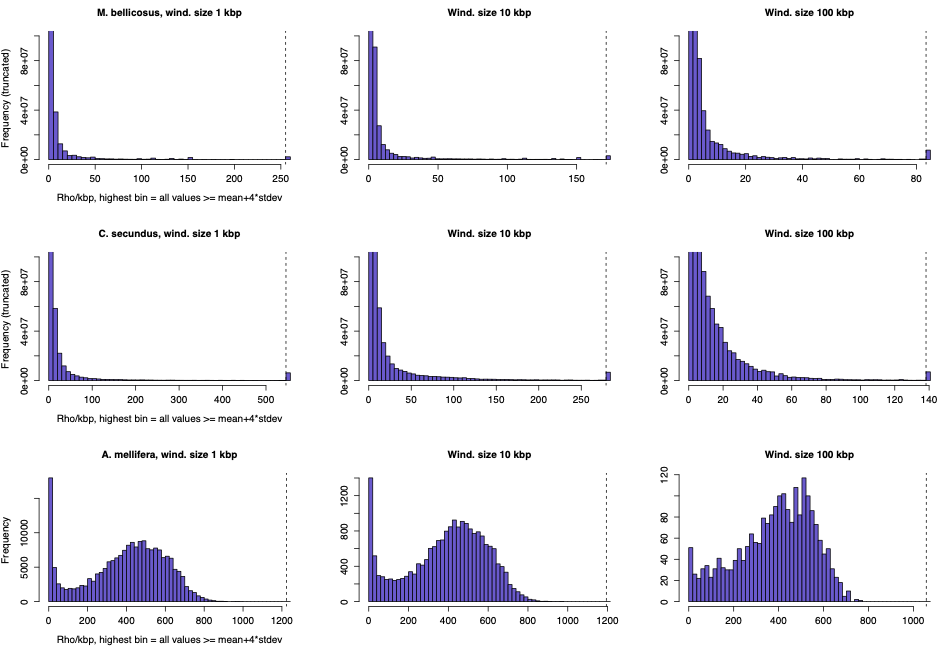

Figure S1. Histograms of *ρ*/kbp for the termites *C. secundus* (top) and *M. bellicosus* (middle) and the honey bee *A. mellifera* (bottom; data from Wallberg et al. 2015), for different window sizes (1 kbp, 10 kbp and 100 kbp). The black dashed line in each histogram marks the value four standard deviations above the mean. All values higher than or equal to this value are included in the highest bin to the right of the dashed line. With 1 kbp windows, the highest bin contains 0.2 % of the *M. bellicosus* values, 0.6 % of the *C. secundus* values and none of the *A. mellifera* values.

a)

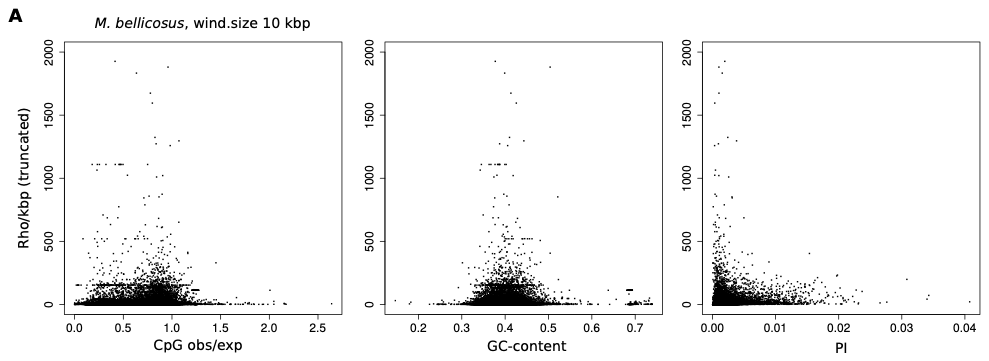

b)

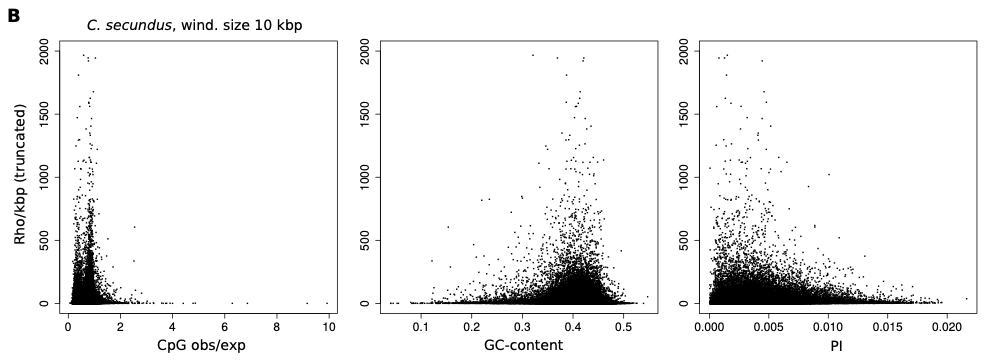

Figure S2. *ρ*/kbp versus CpG_O/E_ (left), *ρ*/kbp versus GC-content (middle) and *ρ*/kbp versus nucleotide diversity (π) (right) in windows of 10 kbp across the genomes of *M. bellicosus* (a) and *C. secundus* (b). Spearman correlations between *ρ*/kbp and the other parameters, weighted by actual window sizes, for *M. bellicosus* and *C. secundus* are 0.24 and 0.097 for CpG; 0.23 and 0.017 for GC-content and 0.48 and 0.31 for nucleotide diversity (π), respectively. Those correlations are all significant with two-tailed p-values < 1 x 10^-4^ calculated by permutation tests with 10,000 iterations.

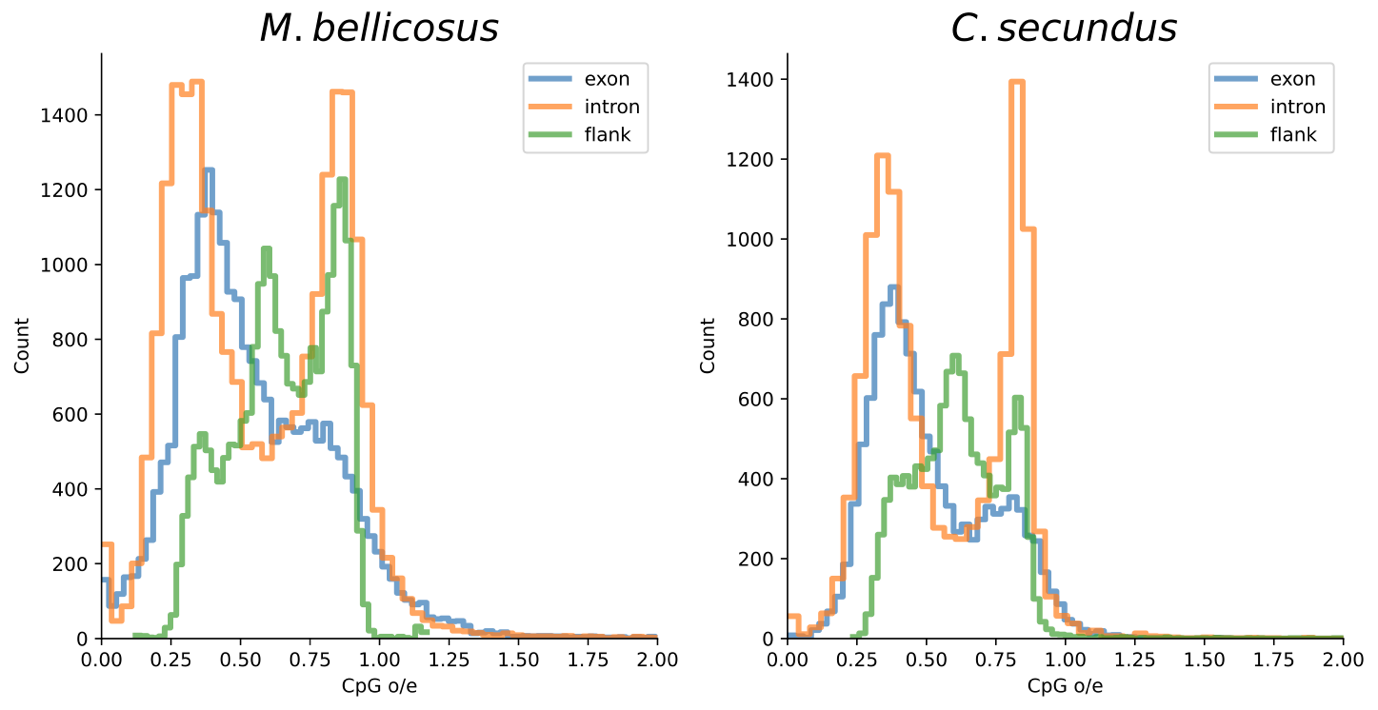

Figure S3. Distribution of CpG_O/E_ in exons, introns and flanking noncoding regions in the *M. bellicosus* and *C. secundus* genomes.

a)

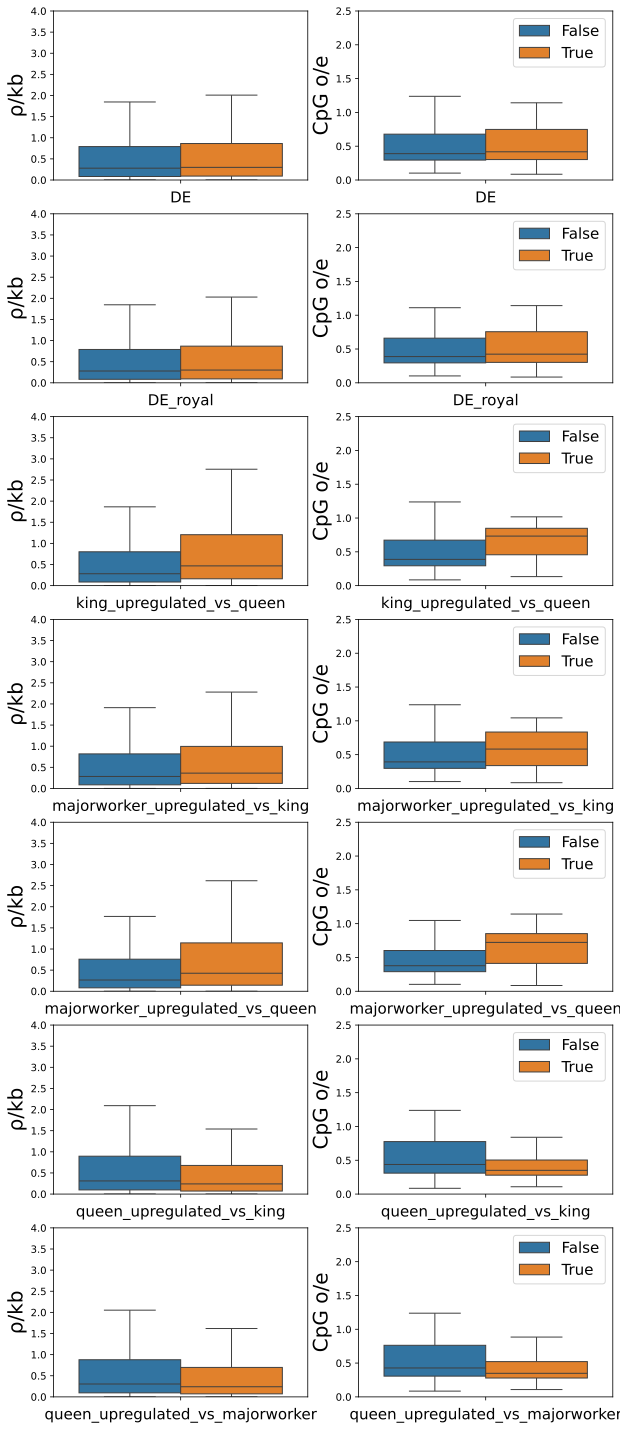

b)

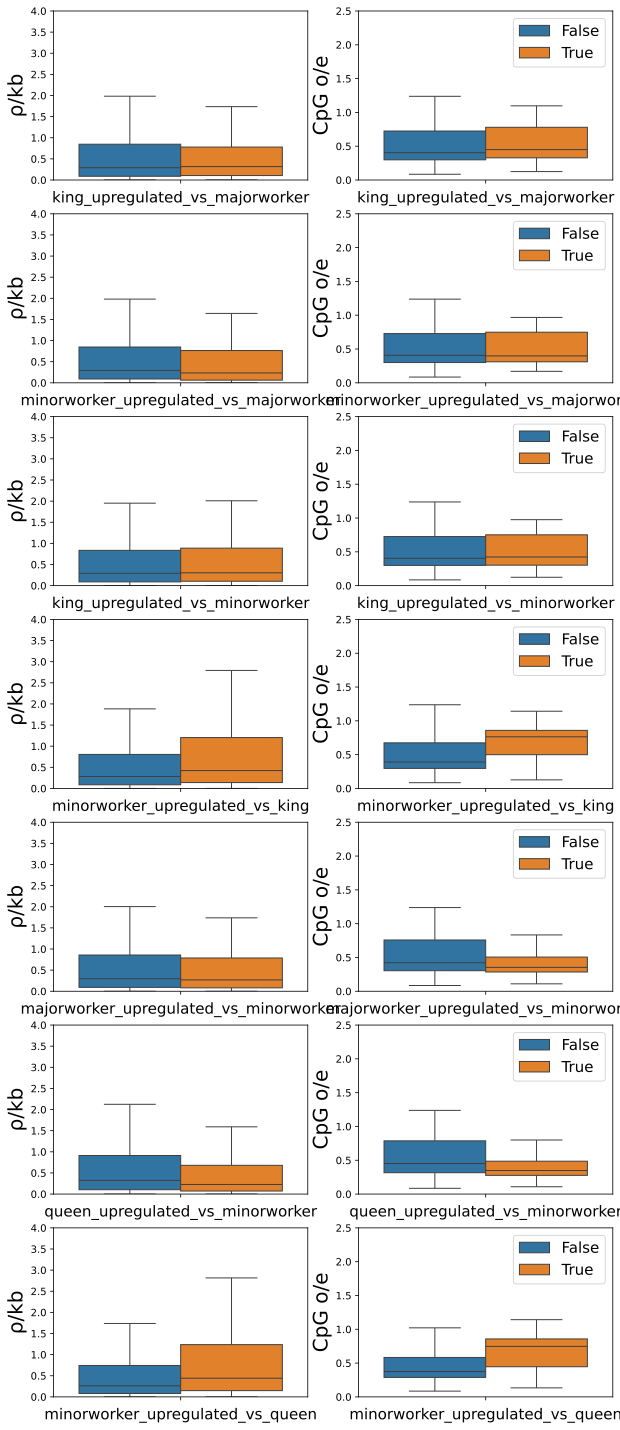

Figure S4. Boxplots of *ρ*/kbp (first column) and CpG_O/E_ (second column) in genes divided by expression categories in *M. bellicosus*.
